# The Pharmacokinetics, Biodistribution and Therapeutic Efficacy of a Trimeric Nanobody Against SARS-CoV-2 in the Golden Syrian Hamster COVID-19 Model

**DOI:** 10.64898/2025.12.17.694727

**Authors:** Parul Sharma, Daniele F Mega, Adam Kirby, Imogen Buckle, Anja Kipar, Jordan J Clark, Eleanor Bentley, Lauren Eyssen, Edyta Kijak, Joanne Herriott, Jo Sharp, Andrew Owen, Miles W Carroll, Raymond J Owens, James P Stewart

## Abstract

Nanobodies are promising medical countermeasures against respiratory viruses, but their local pharmacokinetics and therapeutic window remain incompletely defined. We evaluated the pharmacokinetics, biodistribution and efficacy of a SARS-CoV-2-neutralising trimeric nanobody (A8) in golden Syrian hamsters. Following intranasal administration, A8 concentrations were quantified in nasal wash, bronchoalveolar lavage, lung and serum, and tissue distribution was assessed immunohistologically. Therapeutic efficacy was examined after intranasal treatment before or following challenge with SARS-CoV-2 B.1.351. Intranasal administration of A8 produced sustained respiratory exposure with minimal systemic absorption, whereas intraperitoneal dosing generated transient serum exposure and limited pulmonary delivery. A8 was detected within airway lining material, respiratory epithelial cells, type II pneumocytes and alveolar macrophages. Prophylactic administration 2 h before infection prevented weight loss, while treatment 24 h post-infection accelerated recovery and reduced viral subgenomic RNA in respiratory tissues. Doses as low as 0.1 mg/kg were effective when administered at 24 h post-infection, with no clear dose–response relationship. Treatment delayed until 48 h post-infection was less effective. These findings demonstrate that intranasal A8 achieves favourable compartmentalised exposure and that therapeutic efficacy depends principally on early respiratory delivery rather than dose magnitude, supporting further development of inhaled nanobody therapeutics against SARS-CoV-2.

## 1. Introduction

The coronavirus disease 2019 (COVID-19) pandemic, caused by severe acute respiratory syndrome coronavirus 2 (SARS-CoV-2), has caused over 779 million cases and seven million deaths as of 2026, although from March 2024 onwards many countries stopped reporting (https://data.who.int/dashboards/covid19/). Its emergence has driven an unprecedented global effort to develop effective prophylactic and therapeutic countermeasures. Despite the widespread rollout of vaccines, the continued emergence of antigenically distinct variants of concern, together with waning immunity and the persistence of breakthrough infections, has sustained the need for effective therapeutic agents that can be rapidly deployed against both existing and future SARS-CoV-2 variants [1–4]. Neutralizing-antibody-based treatments, including convalescent plasma, intravenous immunoglobulin, and monoclonal antibodies directed against the viral spike protein, have provided a broad platform for the prevention and treatment of acute SARS-CoV-2 infection, although rapid viral evolution has repeatedly eroded the efficacy of many first-generation monoclonal antibody therapeutics [1, 2, 5].

The receptor-binding domain (RBD) of the SARS-CoV-2 spike (S) protein mediates viral attachment to the host angiotensin-converting enzyme 2 (ACE2) receptor [6–8]. It represents the principal target of neutralizing antibodies elicited by either natural infection or vaccination [9]. Neutralizing antibodies (NAbs) provide an effective and evidence-based intervention during early stages of SARS-CoV-2 infection, and monoclonal antibodies against S have been commonly employed as a therapeutic strategy to neutralize the virus [8, 10]. However, the emergence of Omicron sub lineages has limited the effectiveness of all approved monoclonal antibodies as reported in recent studies [11, 12].

Nanobodies, often referred to as single domain-based variable heavy-chain antibodies (VHHs), are antibody fragments derived from only heavy-chain IgG antibodies found in the *Camelidae* family. They were first discovered in the early 1990s [13] and since then have been extensively reviewed [14–18] Interestingly, the VHH domain can perform antigen-binding independently. Due to their small size (12-15kDa), simple structure, high antigen binding affinity, lower production costs, faster manufacturing and remarkable stability in extreme conditions, nanobodies are an excellent alternative to conventional monoclonal antibodies [2, 18–21]. The majority of anti-SARS-CoV-2 nanobodies identified through phage or ribosome display screening bind the RBD and block ACE2 engagement, and many retain potent, often picomolar-range, neutralizing activity that can be further enhanced through affinity maturation or multivalent (dimeric or trimeric) engineering to increase avidity and cross-linking of adjacent spike trimers [18, 22–24]. Multivalent nanobody constructs in particular have demonstrated superior neutralization potency and resistance to viral escape compared with monomeric formats, and several trimeric nanobodies have shown efficacy *in vivo* when delivered directly to the respiratory tract [18, 23, 24].

Preclinical studies have shown that unmodified nanobodies given intravenously (IV) or intraperitoneally (IN) to animals are cleared extremely rapidly, primarily through glomerular filtration and renal uptake, unless engineered with half-life-extension strategies such as albumin binding, PEGylation, Fc fusion, or multimerization. [25, 26]. However, A key advantage of nanobody-based therapeutics over conventional antibodies is their compatibility with intranasal (IN) or inhaled delivery. Non-invasive respiratory administration enables direct deposition of the therapeutic at the primary site of SARS-CoV-2 entry and replication, the nasal and pulmonary epithelium [2]. Nanobodies are roughly one-tenth the size of conventional IgG antibodies (∼12-15 kDa vs. ∼150 kDa), and this small size, combined with their simple single-domain structure, high thermal stability, and solubility, allows them to be aerosolized without losing structural integrity or neutralizing activity [27, 28]. They show a distinctive pharmacokinetic pattern dominated by lung deposition, local absorption into the bloodstream, and rapid systemic clearance once absorbed. This is a profile well suited to localized respiratory disease treatment with minimal systemic exposure [29].

In our previous studies we have demonstrated the therapeutic efficacy of trimeric nanobodies in the golden Syrian Hamster model of COVID-19 challenged with either the early VIC01 (Pango B) or later Omicron (BA.5) strains of SARS-CoV-2. [18, 30]. However, these studies did not address the biodistribution or kinetics of the nanobodies *in vivo*. In this study, we determined the exposure and efficacy to a trimeric nanobody in the SARS-CoV-2 hamster challenge model.

## Results

### The pharmacokinetics and biodistribution of A8 trimeric nanobody

Our previous studies have shown the effectiveness of trimeric nanobodies in combatting SARS-CoV-2 infection *in vivo*. To evaluate their pharmacokinetics (PK) and efficacy in more detail, we used a nanobody trimer (A8) developed previously and which was extremely potent in neutralising SARS-CoV-2 variant Beta (B.1.351) *in vitro*. [30]. To facilitate detection of the A8 trimer in samples and *in vivo*, a version of the A8 trimer with a c-Myc epitope tag added to the carboxy terminus (A8-Myc) was produced. A8-Myc trimer was administered intranasally (2 mg/kg, IN) to three groups of golden Syrian hamsters. The hamsters were then serially euthanized at 2, 8, and 24 h post-administration. At each time point, nasal wash, bronchoalveolar lavage (BAL), lung tissue and serum samples were collected and the amount of Myc-tagged A8 trimer was quantified by ELISA. A comparative analysis was performed using intraperitoneal (IP) administration of A8-Myc trimer.

The results (Fig 1A) showed that, post-IN administration, the level in BAL at 2 h (1757 ng/ml) was significantly higher than in nasal wash (406 ng/ml), lung (95 ng/ml) and serum ( 18 ng/ml) respectively. In BAL and nasal wash the levels were highest at 2 h and decreased over 24 h post-administration (525 ng/ml) as opposed to lung tissue levels which peaked at 8 h post-administration (247 ng/ml). Only a very low level of A8-Myc nanobody was detected in serum samples obtained at early time points after IN dosing (2 and 8 h; peak 19 ng/ml) and was undetectable at 24 h post-administration.

**Figure 1.**
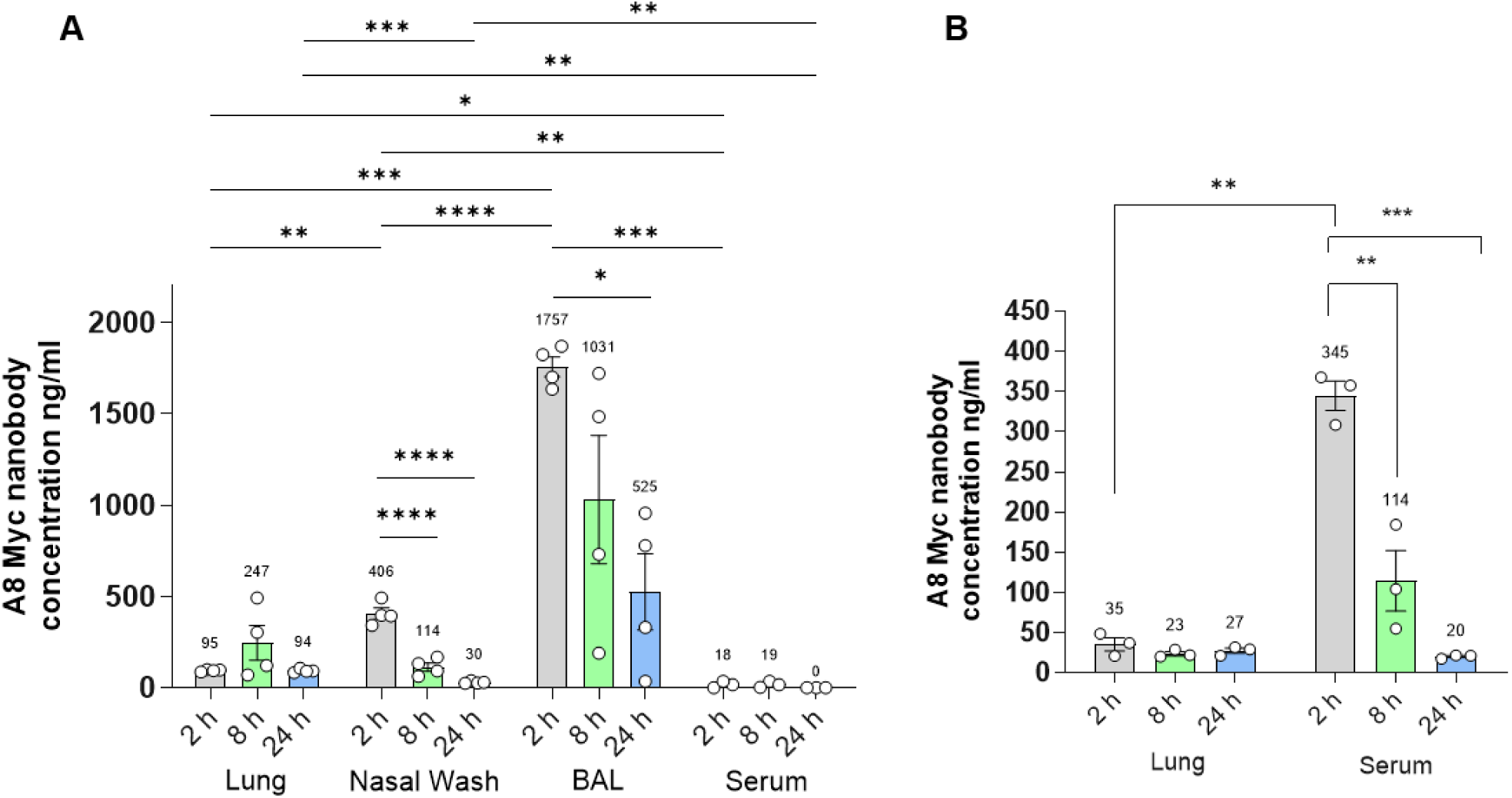
Pharmacokinetic assessment of A8-Myc trimer. Golden Syrian hamsters were dosed with A8 trimer nanobody containing with a Myc tag epitope at the carboxy terminus (A8-Myc) was administered at 2mg/kg intranasally (n=4) **(A)** and intraperitoneally (n=3) **(B).** Hamsters were serially euthanized at 2, 8 and 24 h post dosing to obtain tissue lung, nasal wash, BAL and serum. The A8-Myc nanobody concentration was determined by ELISA. The data represent the mean value ±SEM with the mean value shown above the bars. Comparisons were made using one-way ANOVA with Tukey’s multiple comparisons test. * represents *P ≤ 0.05, **; P ≤ 0.01* ** ; *P ≤ 0.001* and **** ; *P ≤ 0.0001*.

Comparative analysis of the pharmacokinetic profile following IP administration of A8-Myc (Fig. 1B) revealed that the highest serum concentration (344 ng/mL) was observed at 2 h post-administration, followed by a decline to 20 ng/ml over 24 h. Only very low levels (approx. 30 ng/ml) of A8-Myc were seen in lung samples.

To complement the ELISA results, an immunohistological study to detect A8-Myc *in situ* was performed on respiratory tissues of the hamsters dosed via the IN route. At 2 h, A8-Myc was detected cell free in the nasal cavity (in the nasal lining fluid) and attached to the epithelial lining, i.e. the cilia (Fig. 2A, B). It was also detected in the cytoplasm of individual cells in the nasal epithelium including the olfactory epithelium (both sustenticular and olfactory cells; Fig. 2A, B). In the lungs, patches of alveoli contained A8-Myc cell-free in the lumen (Fig. 2C); however, there were also individual alveolar cells, morphologically consistent with type II pneumocytes/alveolar macrophages, that contained A8-Myc in the cytoplasm (Fig. 2C, D). In addition, a variably intense serum reaction was observed in both tissues. At 8 h, staining was less extensive but still obvious in the mucus lining the nasal epithelium and in some epithelial cells (Fig. 2E) as well as free in alveolar lumina and in individual alveolar cells (type II pneumocytes/alveolar macrophages) (Fig. 2F). The serum reaction was still present. There was no evidence of A8-Myc presence in the nose at 24 h post-administration while there was very limited evidence of it in a few alveolar cells. After IP administration, A8-Myc was detected in the serum, mainly in the lung (Supplemental Fig. S3) without evidence of A8-Myc in epithelial cells or in the nasal mucosa or lumen. Staining results in individual animals are presented in Supplemental Table S1.

**Figure 2.**
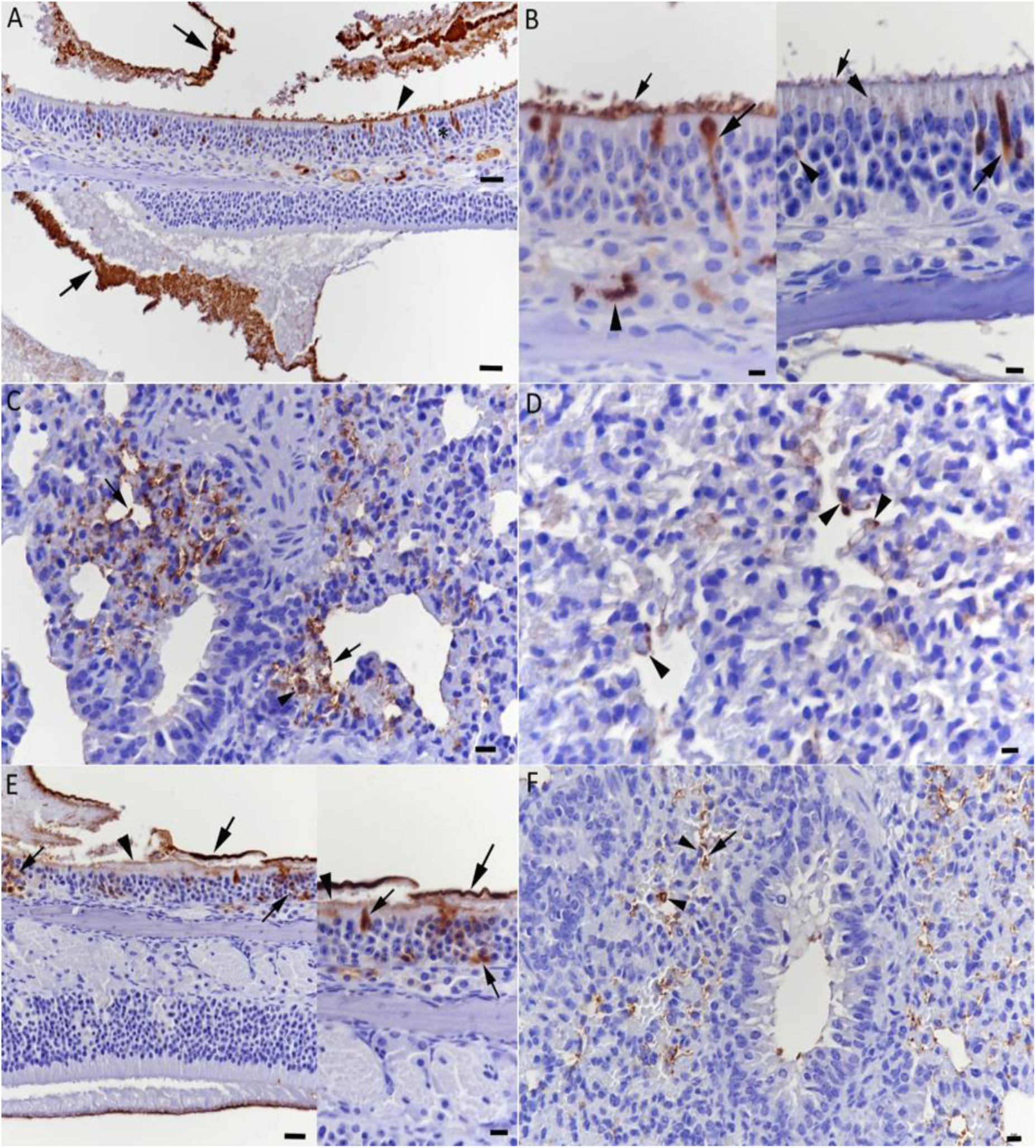
Detection of A8-Myc nanobody in the nose and lung of hamsters. Hamsters were inoculated intranasally with A8-Myc. FFPE tissue sections (2 – 4 µm thick) were stained for the presence of the Myc tag (brown) and counterstained with haematoxylin. **A-D**. 2 h post inoculation. **A, B.** Nose. **A.** Overview from the nasal cavity with lining mucosa. A strong positive reaction is seen as a precipitate in the lumen (arrows) and attached to the luminal surface of the epithelial cells (top image: arrowhead). **B.** Closer views of the nasal mucosa. A strong reaction is seen in the material (mucus) attached to the cilia at the luminal surface of the epithelial cells (small arrows) and diffuse within the cytoplasm of individual cells morphologically consistent with olfactory cells (large arrows) or as a punctate reaction in the luminal cytoplasm (right image: arrowheads) of (sustenticular) cells. The subepithelial layers also contain occasional positive cells (left image: arrowhead) with the morphology of macrophages. **C, D.** Lung. **C.** A strong reaction is seen in material free in the alveolar lumen (arrows) and in individual cells (arrowhead). **D.** A closer view shows an area with a few cells exhibiting a cytoplasmic reaction (arrowheads) and a morphology consistent with type II pneumocytes/alveolar macrophages. **E, F.** 8 h post inoculation. **E.** Nose, overview (left) and closer view (right). A strong reaction is seen in a layer of material (large arrows; likely mucus) attached to the cilia (arrowheads) of the epithelial cells. A few epithelial cells (small arrows) are also found to contain Myc tag. **F.** Lung. Myc tag is seen in material free in alveolar lumina (arrow) and in individual alveolar cells (arrowheads). Hematoxylin counterstain; bars = 25 µm (A, C, E left) and 10 µm (B, D, E right, F).

### Reduced weight loss in A8 trimer-treated Syrian golden hamsters

In a previous study we showed that the A8 trimer neutralizes the SARS-CoV-2 B.1.351 (Beta) variant (NT90 = 0.66 nM) *in vitro* but was less effective against Omicron sub-variants of concern [30]. To assess the effectiveness of the A8 trimer nanobody, it was administered by the IN route both before and after infection and using different doses of the nanobody as shown in the schematic (Fig. 3). The IN route was chosen as this was the most effective route of dosing as shown in the previous analysis of its PK. To assess the clinical disease progression, animals were monitored and weighed daily for weight loss and clinical signs. The hamsters were culled on day 7. The weight loss data (Fig. 4) showed decreased weight loss throughout the study in hamsters treated with the A8 nanobody compared to untreated hamsters. The animals that received 2 mg/kg of A8 nanobody 2 h before infection exhibited no weight loss, with a significant difference when compared with the untreated group (*P* < 0.001 at day 7). Animals treated with 2 mg/kg at 24 h post-infection (p.i.) showed an initial weight loss but from day 3 p.i., their body weight gradually increased with significantly less weight loss (*P* < 0.05) at day 7 compared to the untreated group. The groups that had received the lower doses of A8 trimer, 0.4 mg/kg and 0.1 mg/kg (24 h p.i.), followed the weight loss trend of the 2 mg/kg group, also showing less weight loss (*P* = 0.05 and not significant respectively) compared to the untreated group. There was little indication of a dose response relationship with treatment at 24 h p.i. with 0.1 mg/kg being as effective as 2 mg/kg. The group treated with 0.4 mg/kg at 48 h p.i. showed an initial weight loss similar to the PBS control but then recovery at day 7 p.i. (not significant) indicating that treatment at 48 h p.i. was less effective at controlling clinical signs.

**Figure 3.**
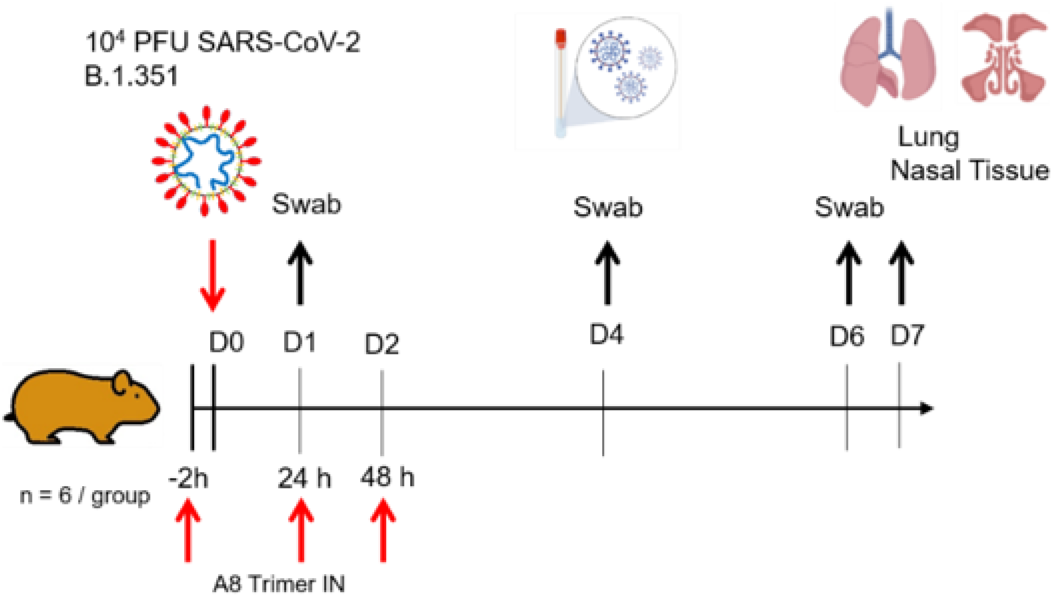
Schematic representation of treatment and challenge infection. Golden Syrian hamsters (n = 6 per group) were infected intranasally with 10^4^ PFU of SARS-CoV-2 strain B.135.1. Individual cohorts were treated intranasally (IN) either 2 h pre-infection or 24 h or 48 h post-infection with 100 μl of A8 trimer at a dose of 2 mg/kg, 0.4 mg/kg or 0.1mg/kg. One group of hamsters received PBS (control group).

**Figure 4.**
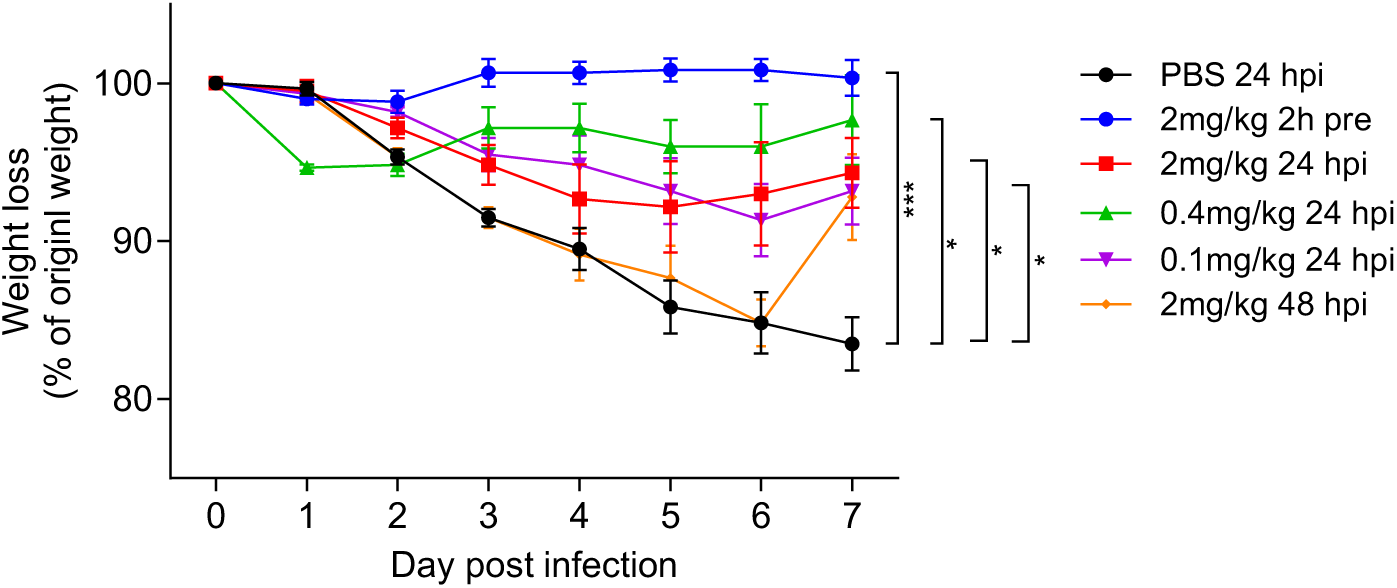
Weight loss after infection and treatment with A8 trimer. Golden Syrian hamsters were challenged intranasally with 10^4^ PFU of SARS-CoV-2 strain B.135.1 and treated with A8 trimer at 2 h pre-, 24 and 48 h post-infection (hpi) as indicated. The data represent mean ±SEM (n=6). Comparisons were made using a repeated-measures two-way ANOVA with Šídák’s multiple comparisons test. *, *P* < 0.05; ***, *P* < 0.001.

### Reduced viral subgenomic load after A8-trimer treatment

Evaluation of the sub-genomic E (SgE) load by RT-qPCR as a surrogate for virus replication was performed on oropharyngeal swabs at 1,4 and 6 days p.i., and nasal turbinate and lung tissue samples on day 7 p.i.. The SgE loads in oropharyngeal swabs collected on day 1 p.i. showed a significant decrease compared with PBS control in all treatment groups apart from the treatment at 2 mg/kg 48 h p.i. (Fig. 5A). At days 4 and 6 p.i. there was a significant decrease in load only with mice receiving 2mg/kg delivered 2 h pre-infection at day 6.

**Figure 5.**
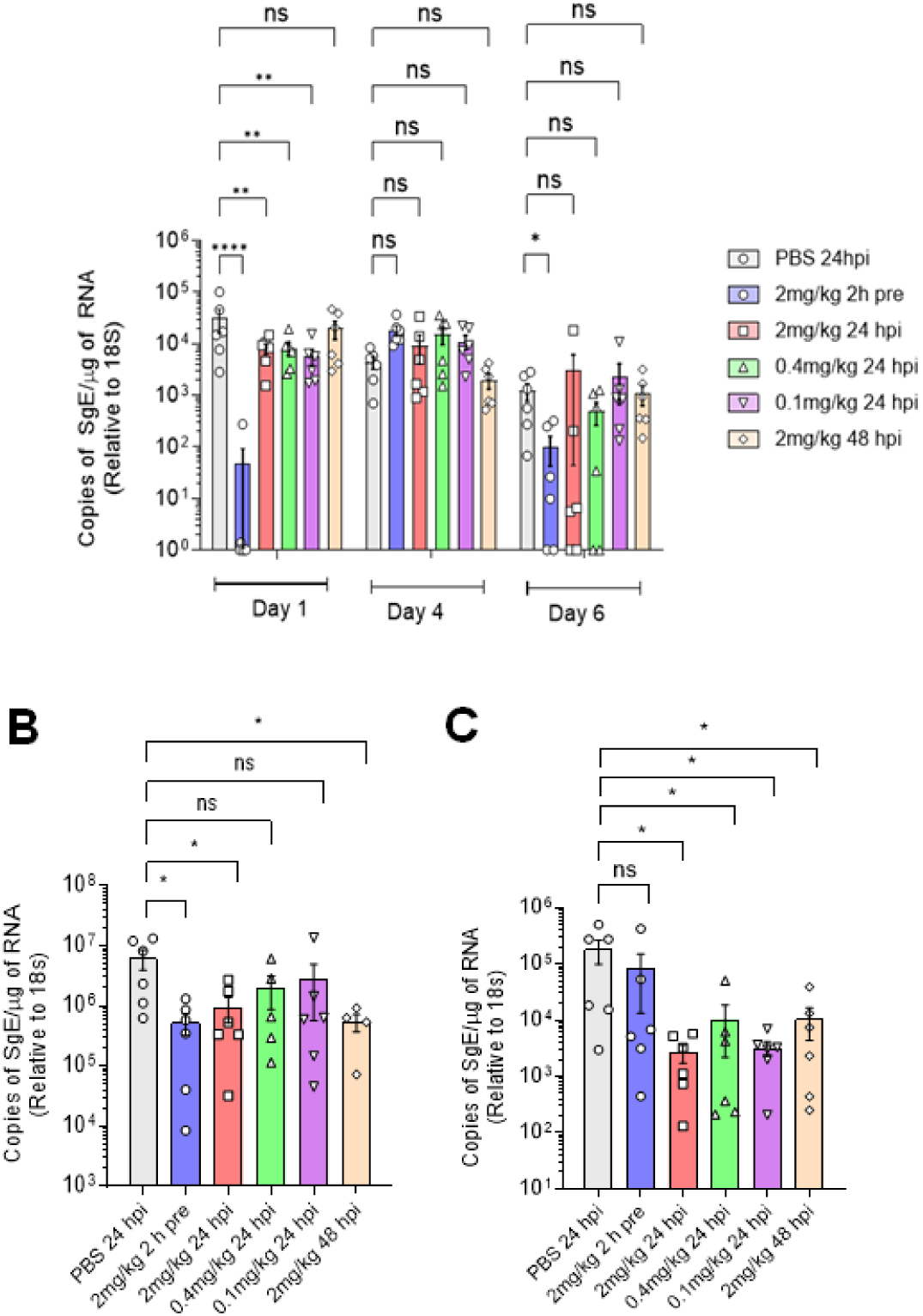
SARS-CoV-2 sgRNA levels in infected hamsters. Hamsters were dosed intranasally with A8 trimer (or PBS) at 2 h pre-, 24 and 48 h post-infection (hpi) as indicated and infected with 10^4^ PFU of SARS-CoV-2 strain B.1.135 in 100 µl PBS intranasally. On days 1, 4 and 6 post-infection oropharyngeal swabs were taken. On day 7 post-infection, all hamsters were euthanised and for subgenomic E RNA (sgE) levels determined by qRT-PCR. Panel **(A)** opharyngeal swabs, **(B)** nasal turbinates **(C)** and lung tissue. Circles represent individual data points and columns mean ±SEM (n=6). Comparisons were made using a Mann-Whitney *U* test. *\*\*, P ≤ 0.01; ****, P < 0.0001*.

The quantification of SgE RNA in nasal tissue (Fig. 5B) revealed a reduction in load in the treated groups compared to the untreated control group. This was significant in the groups given 2 mg/kg 2 h pre-infection, 24 h p.i. and 48 h p.i.. In lung tissue, A8 decreased sgE load significantly when given at all doses as therapeutic 24 h and 48 h p.i. but not when administered 2 h pre-infection. (Fig. 5C). Comparison of different doses of A8 given at 24 h p.i. did not show a difference in effectiveness.

### Histological features and viral antigen expression after A8 trimer treatment

Histological examination of the nasal turbinates and lungs from hamsters euthanized at 7 d p.i. revealed similar changes in all groups (Figs. 6 and Suppl. Fig. S4). Detailed descriptions of all histological changes seen are provided in Supplementary Table S2. Untreated hamsters showed mild multifocal mononuclear infiltration and focal areas with flattened epithelium in the nasal mucosa, often with some mucus and debris in the lumen (Fig. S4). Immunohistology revealed only rare infected epithelial cells. Similar observations were made in hamsters that had received the A8 trimer post infection. In animals that had received A8 trimer 2 h pre-infection, more extensive nasal infection was seen, represented by patches and individual intact, virus antigen positive epithelial cells and sloughed off infected epithelial cells in the lumen. This was accompanied by moderate multifocal to diffuse mononuclear mucosal infiltration, with focal neutrophil infiltrates (Fig. S4B). Consistent with previous reports [18, 30, 31], the lungs of the untreated hamsters exhibited multifocal consolidated areas of pneumonia, with activated and hyperplastic type II cells and infiltration by macrophages and neutrophils, occasionally with a few syncytial cells and degenerate pneumocytes, and combined with mononuclear (peri)vascular infiltrates. In most animals (4/6) this was accompanied by focal bronchiolar epithelial cell hyperplasia (Fig. 6A), providing evidence of regenerative attempts [32]. Viral antigen expression was mainly restricted to a few macrophages within inflammatory infiltrates. However, occasional patches of intact alveoli with infected type I and II pneumocytes were also seen (Fig. 6A). In hamsters that had received the A8 trimer post infection, the histological changes were similar (Fig. 6B-F), with regenerative processes (i.e. bronchiolar epithelial hyperplasia) present in 15/18 animals. The latter was not observed in any hamster that had received A8 trimer 2 h prior to infection. Instead, the histological changes indicated more recent infection, with immunohistological evidence of bronchiolar epithelial cell infection in 4 of the 6 animals, and with areas exhibiting alveoli that contained desquamated epithelial cells and/or alveolar macrophages (Fig. 6B). The extent of the changes and in particular the area affected by inflammatory changes and consolidation varied between groups and individual animals. However, the morphometric analysis did not reveal significant quantitative differences in the amount of free airspace, as an indicator of the amount of functional pulmonary tissue, between the treated groups and the control animals (Fig. 7). Individual animal data are provided in Supplemental Table S1. Taken together, the results indicate that the pre-infection A8 trimer treatment delayed lung infection and thereby also the pulmonary degenerative, inflammatory and regenerative processes.

**Figure 6.**
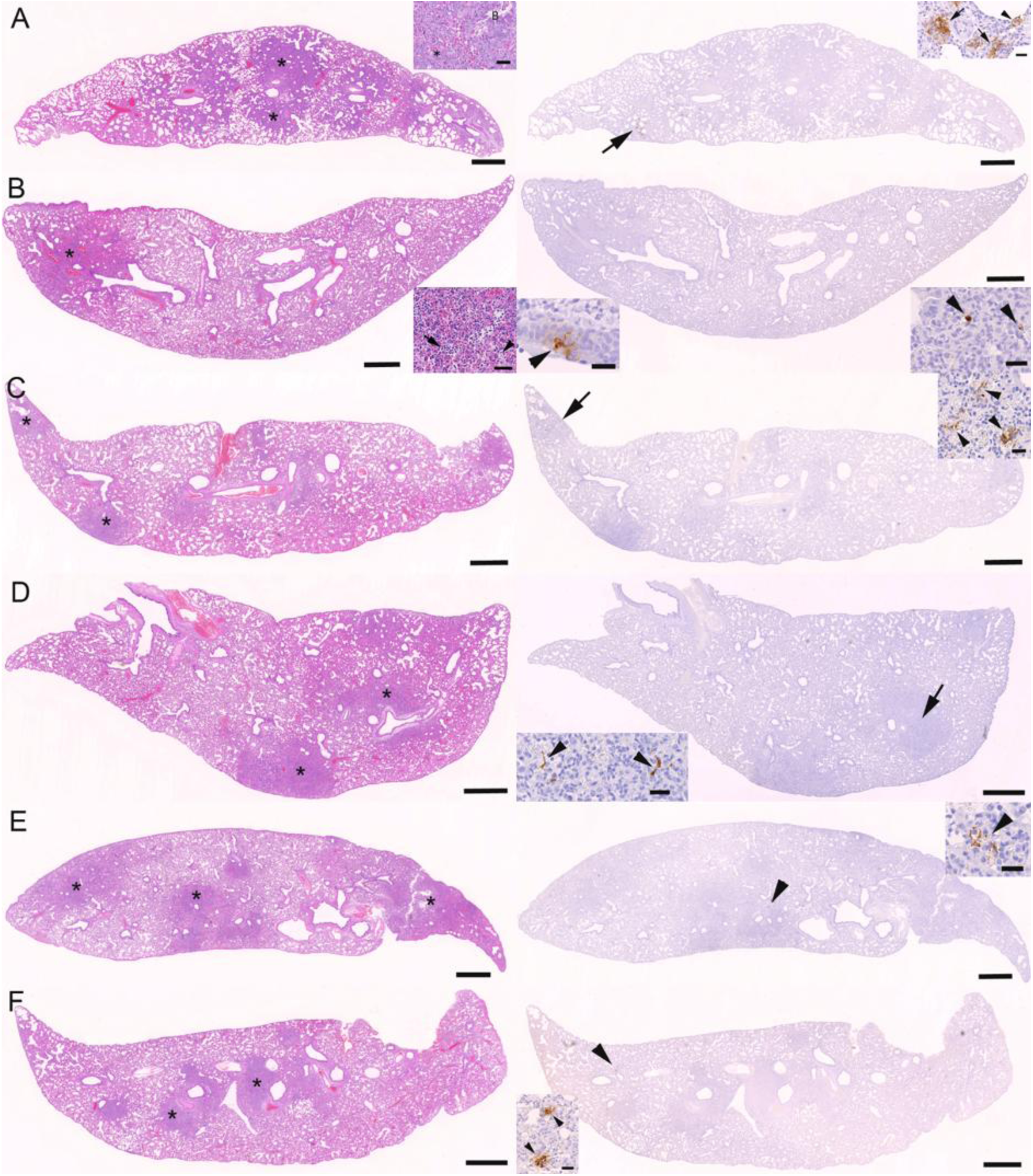
Histological changes and viral antigen expression in the lungs of hamsters. Hamsters were treated with A8 nanobody, infected intranasally with 10^4^ PFU SARS-CoV-2 B.1.351 and euthanised at 7 d p.i. FFPE tissue sections (2 – 4 µm thick) were stained with haematoxylin and eosin (HE; right column) or for SARS-CoV N (brown) and counterstained with haematoxylin (right column). The arrows/arrowheads in the overview images on the right indicate the area shown in the insets. All lungs show areas of parenchymal consolidation (asterisks; left column: HE) and limited expression of viral N (right column). **A.** Animal treated with PBS at 24 h p.i. The inset (HE stain, left image) highlights an area with bronchiolar epithelial hyperplasia (asterisk), representing regenerative attempts. Staining for viral antigen (right image) shows patches of alveoli with infected, SARS-CoV-2 N positive type I and II pneumocytes (inset: arrows) and macrophages (inset: arrowhead). **B.** Animal treated with A8 trimer (2 mg/kg) at 2 h pre-infection. The inset (HE stain, left image) highlights an area with alveoli containing desquamed epithelial cells/alveolar macrophages (arrow) together with proteinaceous fluid (arrowhead) in the lumen. Viral antigen expression (right image) is seen in a few individual bronchiolar epithelial cells (left inset: arrowhead) and rare individual macrophages within inflammatory infiltrates in consolidated areas (right inset: arrowheads). **C.** Animal treated with A8 trimer (2 mg/kg) at 24 h p.i. Viral antigen expression is observed in a focal consolidated area, in pneumocytes (inset: arrowheads). **D.** Animal treated with A8 trimer (0.4 mg/kg) at 24 h p.i. The consolidated area contains a few infected, viral N-positive macrophages (inset: arrowheads). **E.** Animal treated with A8 trimer (0.1 mg/kg) at 24 h p.i. There is very limited viral antigen expression, in a patch of pneumocytes. **F.** Animal treated with A8 (2 mg/kg) at 48 h p.i. Viral N expression is seen in a few patches of pneumocytes (inset: arrowheads). Bars represent 1 mm and 25 µm (insets).

**Figure 7.**
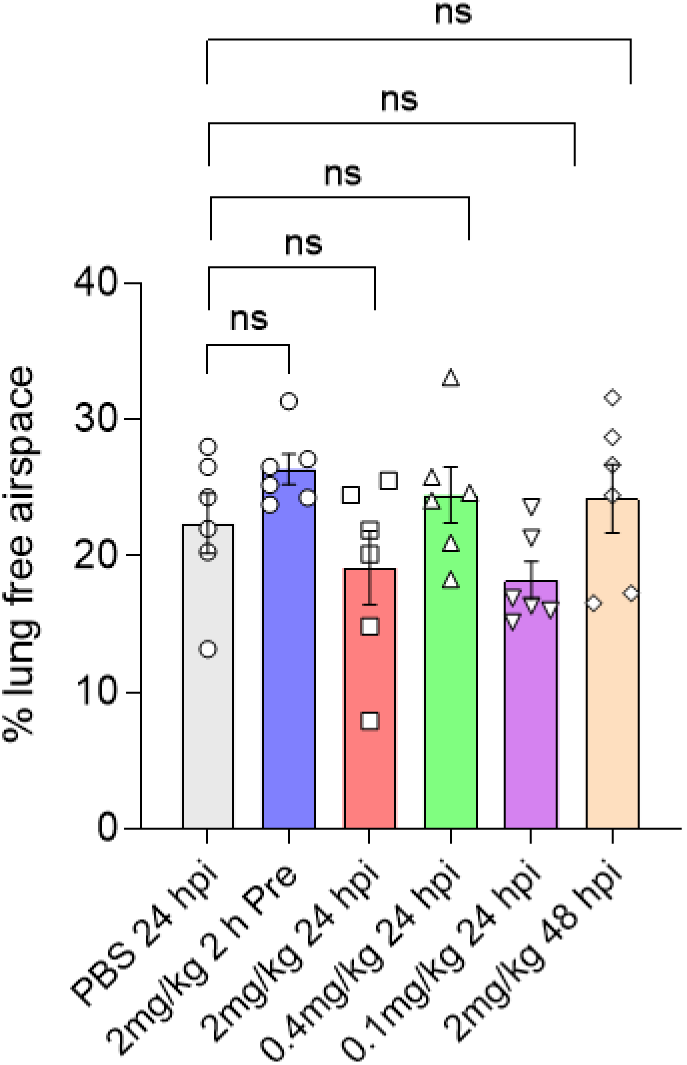
Morphometric analysis of HE-stained sections. Hamsters were treated with A8 nanobody, infected intranasally with 10^4^ PFU SARS-CoV-2 B.1.351 and euthanised at 7 d p.i. FFPE tissue sections (2 –4 µm thick) were stained with haematoxylin and eosin (HE), scanned and analysed using the software programme Visiopharm to quantify the area of non-aerated parenchyma and aerated parenchyma in relation to the total area in the lungs. Comparisons were made using a one-way ANOVA with Dunnett’s test.

## Discussion

This study provides the first integrated assessment of pharmacokinetics, tissue biodistribution, and therapeutic efficacy of a trimeric nanobody. Our previous work had established the *in vitro* neutralizing potency and *in vivo* protective effect of trimeric nanobody constructs without characterizing their exposure profile [8, 30]. The current study uses a trimeric nanobody previously generated against SARS-CoV-2 (A8) in the golden Syrian hamster model [30]. The findings connect two previously separate lines of evidence. Namely, the structural basis of RBD-directed nanobody neutralization and the *in vivo* efficacy of nanobody trimers in hamster challenge models. We show that intranasal (IN) delivery achieves sustained, compartmentalized respiratory tract exposure that correlates directly with protection against weight loss and reduced viral burden.

The pharmacokinetic data show a clear divergence between the two administration routes (Fig. 1). Following IN dosing, A8-Myc concentrations peaked rapidly in bronchoalveolar lavage (BAL) and nasal wash within 2 h, while lung tissue levels peaked later, at 8 hours post-administration, indicating time-dependent redistribution from the airway lumen into the deeper pulmonary compartment. In contrast, intraperitoneal (IP) administration produced a serum concentration peak of at 2 hours that declined by 24 hours, with only minimal nanobody detected in lung tissue at any time point. Only three time points were tested in this study, and the lung shows a non-monotonic pattern which is insufficient for robust non-linear regression or distinguishing between one- and two-compartment kinetics. However, rough approximations indicate elimination half-lives of 54 h in the lung, 13 h in BAL and 6 h in nasal wash after IN inoculation and 5 h in serum after IP inoculation.

This pattern is consistent with previous work. The few pharmacokinetic studies of nanobodies after inhalation indicate high levels in the lung with minimal systemic exposure [33] and that a trimeric nanobody (ALX-0171) against respiratory syncytial virus was found at high local concentrations in BAL fluid and even 24 h after the last dose [29]. The negligible serum exposure detected after IN dosing (undetectable by 24 h) in the current study confirms minimal systemic absorption via the respiratory mucosa, reinforcing that direct airway deposition, rather than systemic distribution, is the primary determinant of A8 trimer availability at the site of viral replication.

In contrast, unmodified nanobodies after systemic administration are rapidly cleared serum half-lives typically measured in hours. This is due to their molecular weight (12–15 kDa) falling well below the ∼60 kDa glomerular filtration threshold and they are cleared by the kidneys. The half-life can be extended by engineering strategies such as albumin-binding domains, Fc fusion, or PEGylation [25, 34]. Studies using radiolabelled or albumin-binding nanobody constructs have shown that fusion to an anti-albumin domain can extend half-life from 1–2 hours to several days by exploiting FcRn-mediated recycling, underscoring that the rapid systemic decline observed for A8 after IP dosing reflects an intrinsic property of the unmodified trimer rather than an assay artifact.

This route-dependence mirrors findings from other nanobody-based SARS-CoV-2 therapeutics. The Pittsburgh inhalable Nanobody 21 (PiN-21) achieved predominant neutralizing activity in BAL fluid, tracheal aspirate, larynx, and nasal wash following nebulized delivery, with dose minimization to 0.2 mg/kg via aerosol compared with 0.6 mg/kg via direct intranasal instillation [35]. Similarly, the nebulized RBD-specific nanobody NIH-CoVnb-112 demonstrated that direct respiratory deposition, rather than systemic circulation, drove protective efficacy in the hamster model [36]. Together with the current A8 findings, these studies establish IN or inhaled delivery as the pharmacologically rational route for nanobody-based respiratory antivirals, sparing the need for the half-life-extension engineering typically required for systemically administered nanobody therapeutics [37].

The immunohistological data shown here (Fig. 2) add spatial resolution to the ELISA-based quantification. A8-Myc was detected both free within the nasal cavity lumen and mucus layer, and internalised within olfactory sustentacular and olfactory sensory cells, as well as within alveolar type II pneumocytes and macrophages. This dual distribution, extracellular reservoir plus intracellular uptake, suggests that the nanobody is not merely transiently present but may be retained within respiratory epithelial and immune cell populations, potentially prolonging local antiviral activity beyond what serum or lavage concentrations alone would suggest.

Previous studies have shown that the *in vitro* neutralisation threshold (NT_90_) for the A8 trimer against the SARS-CoV-2 Beta variant is 0.66 nM, equivalent to approximately 30 ng/mL [30]. Thus, after IN inoculation with 2 mg/ml, the concentration of A8 trimer is at or above the NT_90_ for 24 h in all compartments where SARS-CoV-2 infection occurs (lung (94 ng/ml), nasal wash (30 ng/ml) and BAL (525 ng/ml)). Whereas, after IP inoculation, concentrations of A8 trimer in lung tissue only achieved NT_90_ at 2 h post-administration (35 ng/ml) and were below NT_90_ at 24 hours (27 ng/ml). Thus, the exposure over an extended period after treatment via the IN route argues for this as a much more tractable route for delivery in a clinical setting.

The weight loss and viral load data together indicate that the protective effect of the A8 trimer is highly dependent on the timing of administration relative to infection rather than on dose escalation within the tested range. Prophylactic dosing 2 h before challenge completely abrogated weight loss, while treatment at 24 h post-infection produced significant, though less complete, protection, and treatment delayed to 48 h post-infection showed only a non-significant trend toward recovery. Notably, doses as low as 0.1 mg/kg administered at 24 h post-infection performed comparably to the 2 mg/kg dose, arguing against a straightforward dose-response relationship, at least within this concentration range in this time window.

This pattern is broadly consistent with the dose-sparing efficacy reported for other trimeric nanobody constructs in the hamster model of COVID-19. In our previous studies, the C5 trimer achieved potent therapeutic efficacy at a single IN dose of just 0.4 mg/kg (approximately 40 μg per animal) given 24 hours post-challenge [18], while PiN-21 demonstrated protective efficacy at ultra-low doses down to 0.2 mg/kg by aerosol [35]. The convergence of these independent studies on low effective doses across different nanobody scaffolds supports the broader principle that high intrinsic neutralizing potency, rather than dose magnitude, governs *in vivo* efficacy once a nanobody trimer is delivered directly to the respiratory tract.

The time-dependence of administration observed for A8 trimer was corroborated by the subgenomic E (SgE) RNA data, which showed the most consistent viral load reductions in oropharyngeal swabs at day 1 p.i., and by the finding that A8 trimer significantly reduced lung SgE load when given therapeutically at 24 and 48 h p.i. but not when given prophylactically 2 hours before challenge. This last observation initially appears paradoxical given the clear weight-loss protection conferred by pre-infection dosing, but is explained by the histological findings: animals treated 2 h pre-infection showed no evidence of the bronchiolar epithelial regeneration seen in later-treated groups, and instead displayed histological features consistent with a more recently established infection. Together, these results suggest that prophylactic A8 trimer treatment delayed to some extent, rather than fully prevented, the establishment of pulmonary infection, shifting the disease course later without eliminating detectable viral replication in the lung by day 7. A related pattern was reported for PiN-21, where nebulized therapeutic delivery near the time of infection produced up to a six-log reduction in lung viral titre and rapid reversal of weight loss, suggesting that the degree of viral load reduction achievable is highly sensitive to the precise relationship between dosing time and the kinetics of local viral replication across different nanobody-hamster studies [35]

The absence of a dose-response relationship in the therapeutic dosing window, combined with the demonstrated efficacy of the lowest tested dose (0.1 mg/kg), is consistent with the ultra-low effective doses reported for other inhalable nanobody constructs and suggests that IN A8 trimer dosing regimens could potentially be optimized for lower total protein requirements [38, 39] This would have practical implications for manufacturing cost and scalability, given the comparatively simple and inexpensive production of nanobodies relative to full-length monoclonal antibodies. At the same time, the loss of protective effect when treatment is delayed to 48 hours post-infection defines a relatively narrow post-exposure therapeutic window, a constraint also noted for other nanobody constructs delivered at later time points relative to infection, and this will need to be considered in any translational development pathway given that human patients are rarely diagnosed and treated within hours of symptom onset [18, 35].

Collectively, these findings demonstrate that IN administration of the A8 trimeric nanobody achieves favourable, compartmentalised pharmacokinetics with minimal systemic exposure, consistent with the broader literature on respiratory-delivered nanobody therapeutics, and that this exposure is temporally and spatially linked to measurable reductions in clinical disease and viral load [36, 40]. Early intervention, whether prophylactic or immediately post-exposure, appears more important than dose magnitude for therapeutic benefit, echoing the dose-sparing efficacy reported for other trimeric and inhalable nanobody constructs [35]. These results support continued development of IN-delivered nanobody trimers as non-invasive, rapidly deployable countermeasures against SARS-CoV-2, while highlighting the need for variant-adaptive nanobody engineering, potentially incorporating epitope cocktails or conserved S2-directed constructs, and further work to define the practical limits of the post-exposure treatment window [41].

## Materials and Methods

### Production of A8 trimer nanobodies

The design, construction and production of A8 trimer nanobody was described previously [30]. A vector for expressing the c-Myc tagged A8 trimer was constructed by PCR using the A8 trimer expression vector previously constructed [30] and the following primers:

Fwd 5’ GCGTAGCTGAAACCGGCCAGGTGCAGC 3’

Rev 5’ ACCGTCTCCTCACAGATCCTCTTCAGAGATGAGTTTCTGCTCAAACACCATC AC ‘3

The amplified cDNA incorporating a c-Myc epitope at the 3’ end of the trimer was inserted into the vector pOPINTTG by Infusion cloning. A8 trimer with or without a c-Myc-tag protein was produced by transient expression in expi293™ cells and purified from the cell media by a combination of immobilised metal affinity chromatography and gel filtration [42]. Endotoxin-free buffers were used throughout the purification, and the final products were passed through two Proteus NoEndo™ clean-up columns (Generon, Slough, UK) to reduce endotoxin levels to < 0.1 EU/ml. Endotoxin levels were quantified using the Pierce™ LAL Chromogenic Endotoxin Quantitation Kit (Thermofisher Scientific).

Expi293™ cells were transfected with the pOPINTTG vector expressing the Myc tagged A8 trimer and after 48 h the cells were collected by centrifugation. The cell pellets were then fixed in 10% buffered formalin and paraffin wax embedded for immunohistological examination for use as a positive control for A8-Myc staining.

### A8 trimer ELISA

ELISA was standardized and performed to evaluate the concentration of A8 trimer in bronchoalveolar lavage (BAL), nasal wash, lung tissue and serum samples of hamsters used in the pharmacokinetic study. The 96 well plate was coated with 1mg/ml neutravidin (Thermo Scientific, 31000) overnight and washed 5X with PBS and 0.05% Tween20. Furthermore, biotinylated Wuhan RBD antigen (50nM/well) was added and incubated for 1h, at room temperature (RT) and 500 rpm on a vibrating shaking platform. The A8 trimer Myc was used to generate a standard curve after 2-fold serial dilution. BAL, nasal wash, lung tissue supernatants and serum were added in duplicate into a 96 well plate and incubated for 1 h. Anti-Myc-HRP (Invitrogen R951-25) was used at 1:5000 in 0.1% BSA-PBS and finally ABTS substrate (Fisher Scientific, 37615) was used and plate absorbance was measured at 405 nm after 15 min of incubation.

### In vivo studies

All animal work was approved by University of Liverpool Animal Welfare and Ethical Review Body and performed under UK Home Office project licence PP4715265. Golden Syrian hamsters (male, 8 - 10 weeks old) were obtained from Janvier labs (France). Hamsters were kept in an SPF (Specific Pathogen Free) unit in individually ventilated cages and provided *ad libitum* with water and pellet food.

### Pharmacokinetic evaluation of A8 trimer

A8 trimer was administered to hamsters intranasally (n = 4 per group) at 2 mg/kg body weight in 100 µl volume of PBS, and the hamsters were serially sacrificed at 2 h, 8 h and 24 h post-inoculation with a high dose of pentobarbitone. BAL, nasal wash and blood were collected. Separately, A8-Myc trimer was administered to hamsters intraperitoneally (n = 3 per group) at 2 mg/kg body weight in 100 µl volume of PBS. They were serially sacrificed at 2, 8 and 24 h post-administration and lung and serum samples were taken for A8 trimer ELISA. Lungs with heart as well as the heads from all animals were dissected and fixed in 10% buffered formalin for histological and immunohistological examination.

### Animal challenge infection

Hamsters were randomly assigned into multiple cohorts (n=6). They were administered 2 mg/kg of the A8 nanobody intranasally in a 100 µl volume of PBS either 2 h before or 24 h or 48 h after intranasal infection with 1×10^4^ PFU of SARS-CoV-2 B.135.1 in 100 µl of PBS. One group served as an untreated (PBS only) infected control. The weights and health status of the hamsters were monitored daily for weight loss and clinical signs. Oropharyngeal swabs were taken from all groups at days 1, 4 and 6 post-infection. At the conclusion of the experiment (day 7 post-infection), the hamsters were euthanized with a high dose of pentobarbitone and dissected immediately after death. Lung and nasal turbinate tissue samples were collected for further analysis.

### Tissue sample processing for qRT-PCR

The upper right lobe of the lung was homogenised in 1 ml TRIzol^TM^ reagent (Invitrogen) using a Bead Ruptor 24 (Omni International) at 2 meters per second for 30 sec. The homogenate was centrifuged at 12000xg for 5 min, followed by full TRIzol RNA (Invitrogen) extraction as per manufacturer’s instruction. The extracted RNA was re-suspended in 60 µl of nuclease free water and quantified using a Nanodrop (Thermofisher). Genomic DNA contamination was removed using the TURBO DNA-free™ Kit (Thermofisher) as per manufacturer’s instructions.

### Viral RNA quantification

The SARS-CoV-2 subgenomic E (SgE) RNA was quantified as a surrogate of virus replication. The SgE primers and probe, previously described by [43] were used at final concentrations of 400 nM and 200 nM, respectively. Murine 18S rRNA primers and probe were used at the concentrations 400 nM and 200 nM to normalize N1 and SgE qPCR. Standard curves for N1, SgE, and 18S were generated using PCR-amplified fragments obtained with their respective qPCR primers. cDNA synthesis was performed using SuperScript IV Reverse Transcriptase (Thermo Fisher Scientific), and the PCR was carried out using Q5® High-Fidelity 2X Master Mix (New England Biolabs) according to the manufacturer’s instructions.

For generation of qPCR standards, PCR-amplified segments of SARS-CoV-2 SgE, and murine 18S rRNA were cloned into plasmids using the TOPO® TA Cloning Kit (Thermo Fisher Scientific). Plasmid DNA was purified and linearised, using the QIAquick PCR Purification Kit (Qiagen) Purified linearised plasmids were then serially diluted 10-fold from 10^10^ to 10^4^ copies per reaction to generate standard curves for absolute quantification.

### Histology, immunohistology and morphometric analysis

From animals in the PK study, the lungs, heart and entire heads were fixed in 10% buffered formalin for appr. 48 h, then transferred to 70% ethanol until further processing. The heads were deskinned and sawn (coronal section behind the eyes) using a diamond saw (Exakt 300; Exakt) and the nose (including nasal turbinates) gently decalcified in RDF (Biosystems) for twice 5 days, at RT and on a shaker. Afterwards, the nose was sliced (2 mm wide coronal tissue sections), and lungs, hearts and noses routinely paraffin embedded. Consecutive sections (2-4 µm) were prepared and stained with hematoxylin-eosin (HE) for histological examination or subjected to immunohistological staining for Myc-tag, using a mouse anti-Myc-tag antibody (clone 9B11; Cell Signaling Technology) and the horseradish peroxidase (HRP) method. Briefly, after deparaffination, sections underwent antigen retrieval in Tris/EDTA buffer (pH 9) for 20 min at 98 °C, followed by overnight incubation at 4 °C with the primary antibody (diluted 1:2,000 in dilution buffer; Agilent Dako). This was followed by blocking of endogenous peroxidase (peroxidase block, Agilent Dako) for 10 min at RT and incubation with the detection systems (Envision rabbit; Dako Agilent) for 30 min an RT. Incubations were all performed in an autostainer (Dako Agilent). Sections were subsequently counterstained with haematoxylin. A formalin-fixed cell pellets prepared from expi293™ cells transfected with pOPINTTG vector served as positive control. This yielded a granular to diffuse expression pattern in a large proportion of the cells (Supplemental Figure S1A). A non-transfected pellet as well as the tissues from an untreated hamster served as negative controls (Supplemental Figures S1B, S2A and B).

From animals subjected to the SARS-CoV-2 challenge infection, the left lobe of the lung and head was dissected immediately after euthanasia and fixed in 10% neutral buffer formalin for 48 h, then transferred to 70% ethanol until further processing. The lung was longitudinally cut in half and routinely paraffin wax embedded. The nose was processed as for the PKS study. Consecutive sections (3–4 µm) were prepared and stained with HE for histological examination or subjected to immunohistological staining for SARS-CoV-2 nucleoprotein (N), as previously described, using a rabbit anti-SARS-CoV N antibody (Rockland Immunochemicals Inc.) following a previously published protocol [18].

For morphometric analysis, HE-stained sections from the left lungs were scanned using the NanoZoomer-XR C12000 (Hamamatsu, Japan) and analyzed with Visiopharm software (version 2020.08.1.8403; Visiopharm, Denmark) to quantify the areas of aerated and non-aerated parenchyma relative to the total lung parenchyma area, as previously described [18].

## Acknowledgements

This work was supported by Wellcome Trust grant (223733/Z/21/Z) to RJO, JPS, MWC and the Rosalind Franklin Institute funding delivery partner EPSRC. The authors are grateful to the technical team in the Histology Laboratory, Institute of Veterinary Pathology, Vetsuisse Faculty, University of Zurich, for excellent technical support.

