## Supplementary files for "The Pharmacokinetics, Biodistribution and Therapeutic Efficacy of a Trimeric Nanobody Against SARS-CoV-2 in the Golden Syrian Hamster COVID-19 Model"

### Supplementary materials

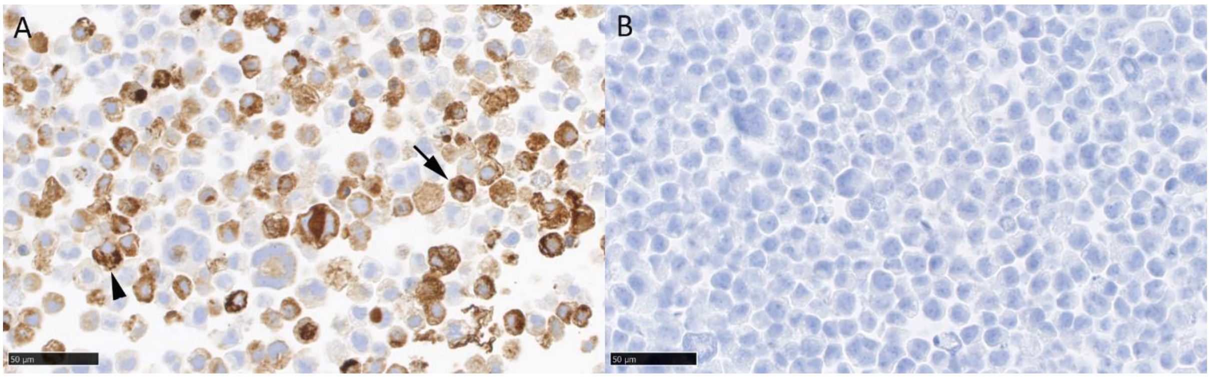

**Supplemental Figure S1**. **Positive and negative control staining for Myc-tag.** Cell pellets prepared from expi293™ cells transfected with pOPINTTG vector were stained by immunohistology for Myc-tag. **A.** A large proportion of cells show a granular (arrowhead) or diffuse (arrow) cytoplasmic reaction, confirming expression of Myc-tag. **B.** The non-transfected cell pellet does not show any reaction. Hematoxylin counterstain; bars represent 50 µm.

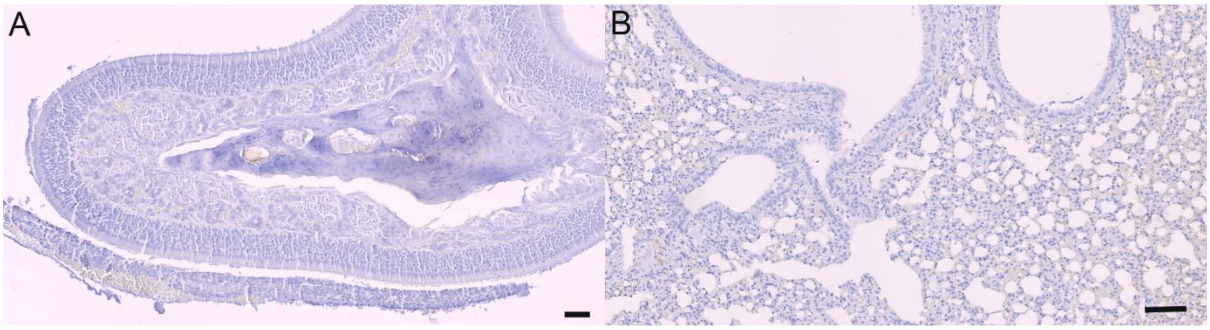

**Supplemental Figure S2**. **Staining for Myc-Tag in untreated hamster tissues**. FFPE tissue sections (2 – 4 µm thick) were stained for Myc-tag (brown) and counterstained with haematoxylin. (A) nose and (B) lung from an untreated hamster, yielding no reaction. Bars represent 50 µm (A) and 100 µm (B).

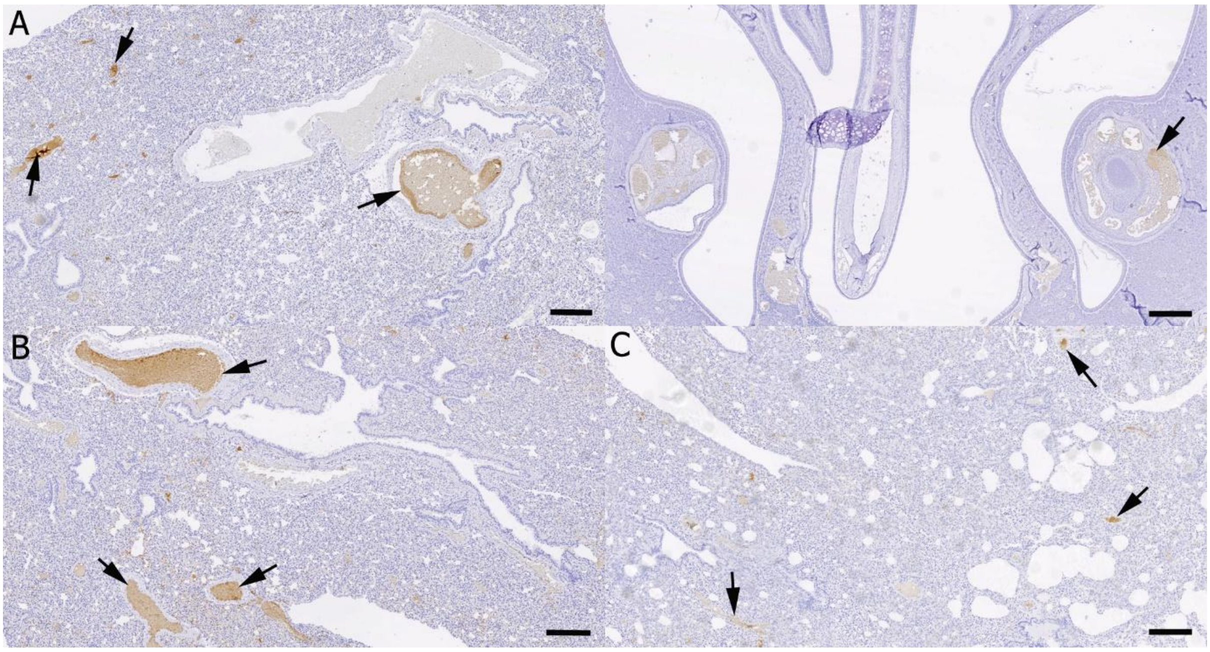

**Supplemental Figure S3**. **Staining for Myc-tag in the nose and lung from hamsters after intraperitoneal administration of A8-Myc. A**. Lung (left) and nose (right) at 2 h post treatment. There is a moderate serum reaction in the lung and a very weak serum reaction in a vessel in the nose (arrows). **B**. Lung at 8 h post treatment, showing a moderate serum reaction (arrows). **C**. Lung at 24 h post treatment, with moderate serum reaction (arrows). Immunohistology, hematoxylin counterstain; bars represent 250 µm.

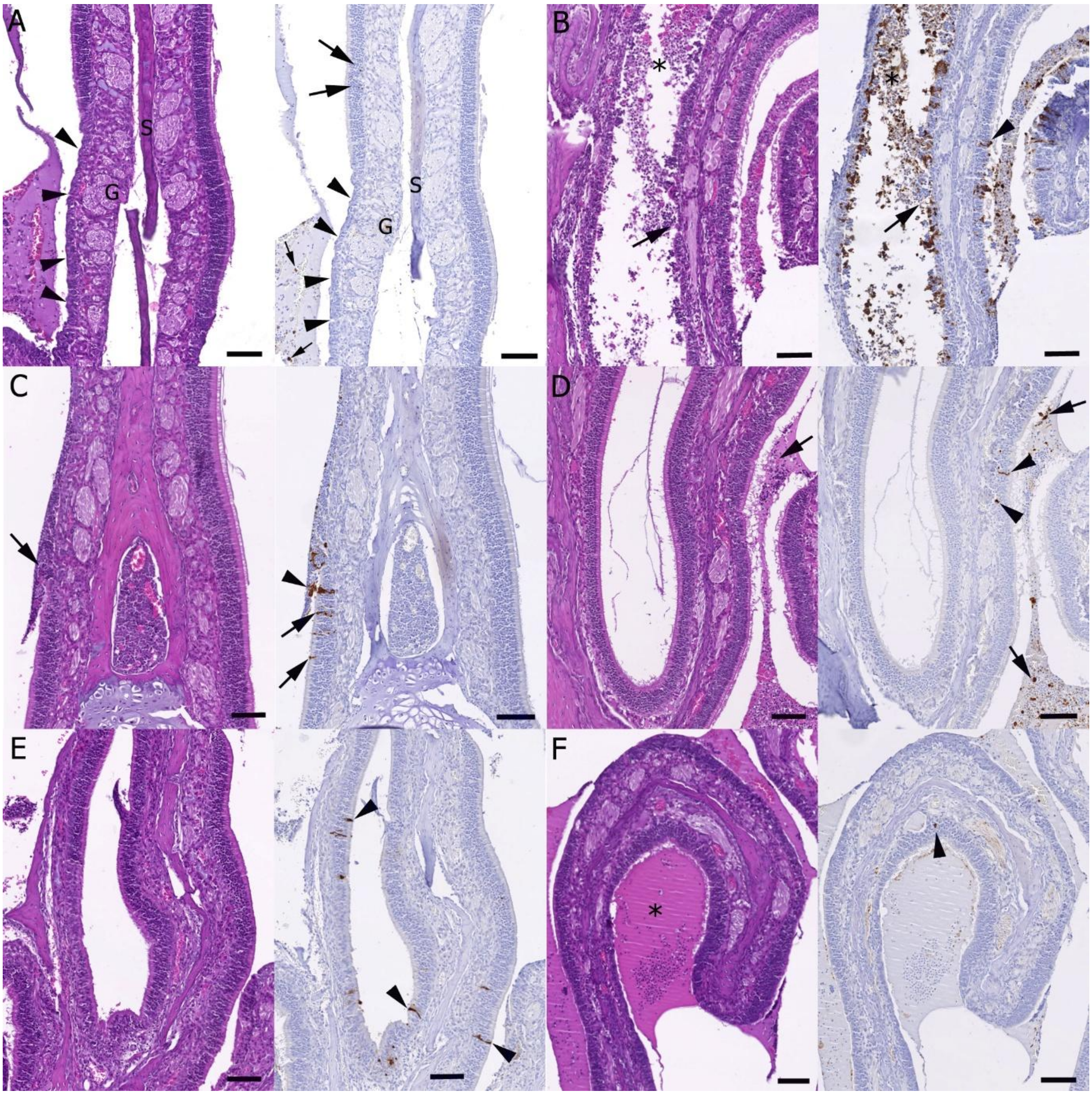

**Supplemental Figure S4**. **Histological changes and viral antigen expression in the nasal turbinates of hamsters.** Hamsters were treated with A8 nanobody, infected intranasally with $10^4$ PFU SARS-CoV-2 B.1.351 and euthanised at 7 dpi. FFPE tissue sections (2 – 4 µm thick) were stained with haematoxylin and eosin (HE; A-F: left) or for SARS-CoV NP (brown) and counterstained with haematoxylin (A-F: right). **A**. Animal treated with PBS at 24 h post infection (hpi). The epithelium is focally flattened (arrowheads), consistent with prior damage. G: subepithelial glands; S: cartilagenous septum. There are individual SARS-CoV-2 NP positive epithelial cells (large arrows) and a few positive degenerate epithelial cells in the mucus filling the lumen. **B**. Animal treated with A8 trimer (2 mg/kg) at 2 h prior to infection. The lumen contains abundant degenerate cells and debris (asterisk), and there is focal loss of the epithelium (arrows). Individual intact epithelial cells are also found to carry viral antigen (arrowhead). **C**. Animal treated with A8 trimer (2 mg/kg) at 24 hpi. There is patch with loss of epithelium (arrow, HE stain) and viral antigen expression in intact (arrows) and degenerate, sloughed off (arrowhead) epithelial cells. **D**. Animal treated with A8 trimer (0.4 mg/kg) at 24 hpi. The epithelium is widely unaltered. Individual intact epithelial cells (arrowheads) as well as a few degenerate cells in debris in the lumen (arrows) express viral NP. **E**. Animal treated with A8 trimer (0.1 mg/kg) at 24 hpi. The epithelium is widely unaltered. Asterisk: mucus with some debris in lumen. Individual intact epithelial cells express viral antigen (arrowheads). **F**. Animal treated with A8 (2 mg/kg) at 48 hpi. The epithelium is widely unaltered. One individual intact epithelial cell is found to express viral antigen (arrowhead). Bars represent 100 µm.

**Supplementary Table S1.** Myc-Tag expression in lung and nasal turbinates (mucosa) of Syrian hamsters after intranasal (IN) or intraperitoneal of 2 mg/kg A8 trimer. Animals were examined 2 h, 8 h or 24 h post administration (hpa). They did not exhibit and any histopathological changes in the examined tissues.

| **Animal No** | **Mode of application; hpa** | **A8-Myc expression (lung, nasal mucosa)** |
|---|---|---|
| A1.1 | IN; 2 hpa | **Lung**: one lobe with reaction lining BEC and mod serum reaction; rare pos type II pn/alveolar macrophage<br>**NT**: strong reaction in nasal lining fluid; mod serum reaction |
| A1.2 | IN; 2 hpa | **Lung**: large focal areas with reaction lining BEC and weak to moderate serum reaction<br>**NT**: strong reaction in nasal lining fluid and focally along NEC, in individual NEC and underlying macrophages; weak to mod serum reaction |
| A1.3 | IN; 2 hpa | **Lung**: focal area with reaction in alveolar lumina and within individual alveolar macrophages; weak to moderate serum reaction<br>**NT**: mod reaction in nasal lining fluid; focal reaction in NEC; mod serum reaction |
| A2.1 | IN; 8 hpa | **Lung**: large area with strong reaction in bronchiolar and alveolar lumina and occ pos type II pn/alveolar macrophage; mod to strong serum reaction<br>**NT**: mod reaction in nasal lining fluid; occ reaction in individual NEC and underlying macrophages; mod serum reaction |
| A2.2 | IN; 8 hpa | **Lung**: patches with reaction along AEC and in occ type II pn/alveolar macrophages; weak to mod serum reaction<br>**NT**: mod reaction in nasal lining fluid; occ patches with a few individual pos NEC; mod serum reaction |
| A2.3 | IN; 8 hpa | **Lung**: patches with reaction along AEC and in occ type II pn/alveolar macrophage; mod serum reaction<br>**NT**: mod reaction in nasal lining fluid; several patches with a few individual pos NE; mod serum reaction |
| A3.1 | IN; 24 hpa | **Lung**: a few small random patches with some reaction in alveoli and individual type II pn/alveolar macrophages; weak serum reaction<br>**NT**: weak serum reaction |
| A3.2 | IN; 24 hpa | **Lung**: a few small random patches with some reaction in alveoli and individual type II pn/alveolar macrophages; weak serum reaction<br>**NT**: weak serum reaction |
| A3.3 | IN; 24 hpa | **Lung**: a few small random patches with some reaction in alveoli and individual type II pn/alveolar macrophages; weak serum reaction<br>**NT**: weak serum reaction |
| B1.1 | IP; 2 hpa | **Lung**: weak serum reaction; **NT**: neg |
| B1.2 | IP; 2 hpa | **Lung**: weak to moderate serum reaction;<br>**NT**: occasional weak serum reaction |
| B1.3 | IP; 2 hpa | **Lungs**: weak to moderate serum reaction; **NT**: neg |
| B2.1 | IP; 8 hpa | **Lung**: mod serum reaction; **NT**: occasional weak serum reaction |
| B2.2 | IP; 8 hpa | **Lung**: weak to mod serum reaction; **NT**: neg |
| B2.3 | IP; 8 hpa | **Lung**: mod serum reaction; **NT**: neg |
| B3.1 | IP; 24 hpa | **Lung**: weak serum reaction; **NT**: neg |
| B3.2 | IP; 24 hpa | **Lung**: weak serum reaction; **NT**: neg |
| B3.3 | IP; 24 hpa | **Lung**: weak serum reaction; **NT**: neg |

**Legend**: AEC – alveolar epithelial cells; BEC – bronchiolar epitethelial cells; hpa – hours post administration; IN – intranasal; IP – intraperitoneal; mod – moderate; NEC – nasal epithelial cells (incl. olfactory epithelial cells); NT – nasal turbinates; occ – occasional; pn - pneumocyte

**Supplementary Table S2.** Relevant histological changes and SARS-CoV-2 nucleoprotein staining in Syrian hamsters after intranasal (IN) infection with $10^4$ PFU SARS-CoV-2 B1.351 and euthanised at 7 days post infection. The control group (Cohort 1; C1) received PBS 24 h post infection (hpi) IN, Cohort 2 (C2) received 2 mg/kg A8 trimer 2 h pre infection, Cohort 3 (C3) received 2 mg/kg A8 trimer 24 hpi, Cohort 4 (C4) received 0.4 mg/kg A8 trimer 24 hpi, Cohort 5 (C5) received 0.1 mg/kg A8 trimer 24 hpi, and Cohort 6 (C6) received 2 mg/kg A8 trimer 48 hpi.

| Animal No | Treatment | Histological changes and viral antigen expression (lung, nasal mucosa) | Virology (PCR)[1] |
|---|---|---|---|
| 1.1 | PBS, 24 hpi | **Lung (HE)**: extensive mf consolidation with activated and hyperplastic type II pn and BEC hyperplasia; focal areas with syncytial cell formation, deg pn, infiltrating macrophages and NL<br>**vAg**: a few small patches of alveoli with pos AEC and macrophages in areas with deg cells; rare individual pos macrophages within inflammatory infiltrates | 2.76e+005 |
| | | **NT (HE)**: very mild mf mononuclear mucosal infiltration; some mucus with debris in lumen<br>**vAg**: EC neg, small amount of cell free vAg in debris in lumen | 1.2e+007 |
| 1.2 | PBS, 24 hpi | **Lung (HE)**: very extensive mf consolidation with activated and hyperplastic type II pn and marked BEC hyperplasia; focal areas with syncytial cell formation, deg pn, infiltrating macrophages and NL and desquamative component; mild mononuclear pv infiltration<br>**vAg**: mf disseminated, often large patches of alveoli with pos AEC outside consolidated areas and within areas with deg cells; occ. individual pos macrophages within inflammatory infiltrates | 2.76e+005 |
| | | **NT (HE)**: mild mf mononuclear mucosal infiltration, focal areas with flattened epithelium; sloughed off EC and debris in lumen<br>**vAg**: disseminated several individual pos intact EC, also desquamed pos EC and cell free viral Ag in debris in lumen | 1.33e+007 |
| 1.3 | PBS, 24 hpi | **Lungs**: very extensive mf consolidation with activated and hyperplastic type II pn, marked BEC hyperplasia, and infiltrating macrophages and NL; mild mononuclear pv infiltration<br>**vAg**: a few small patches of alveoli with pos AEC; low number of pos macrophages within inflammatory infiltrates | 2.96e+003 |
| | | **NT (HE)**: mild mononuclear mucosal infiltration<br>**vAg**: EC neg EC; small amount of cell free vAg in mucus in lumen | 1.12e+006 |
| 1.4 | PBS, 24 hpi | **Lung (HE)**: mf consolidation with activated and hyperplastic type II pn and infiltrating macrophages and NL; mild mononuclear pv infiltration<br>**vAg**: several small patches of alveoli with pos AEC; rare pos macrophages within inflammatory infiltrates | 1.54e+004 |
| | | **NT (HE)**: mild mf mononuclear mucosal infiltration, focal areas with flattened epithelium; sloughed off EC and debris in lumen<br>**vAg**: EC neg, some desquamed pos EC and cell free vAg in debris in lumen | 8.21e+006 |
| 1.5 | PBS, 24 hpi | **Lung (HE)**: mf consolidation with activated and hyperplastic type II pn and infiltrating macrophages and NL; mild mononuclear pv infiltration<br>**vAg**: several small patches of alveoli with pos AEC; rare pos macrophages within inflammatory infiltrates | 1.84e+004 |
| | | **NT (HE)**: mild mf mononuclear mucosal infiltration, focal areas with flattened epithelium; sloughed off EC and debris in lumen<br>**vAg**: EC neg, some desquamed pos EC and cell free vAg in debris in lumen | 2.33e+006 |
| 1.6 | PBS, 24 hpi | **Lung (HE)**: mf consolidation with activated and hyperplastic type II pn and infiltrating macrophages and NL, in some areas with desquamative component; mild mononuclear pv infiltration<br>**vAg**: multiple, often very large patches of alveoli with pos AEC | 5.04e+005 |
| | | **NT (HE)**: mild mf mononuclear mucosal infiltration, focal areas with flattened epithelium; sloughed off EC and debris in lumen<br>**vAg**: disseminated numerous individual and small patches of pos (intact) EC, also desquamed pos EC and cell free vAg in lumen | 6.28e+005 |

| | | | |
|---|---|---|---|
| 2.1 | 2 mg/kg A8 trimer, 2 h pre-infection | **Lung (HE)**: mild pb mononuclear LC dominated infiltration, with occ LC between BEC; one bronchiole with a few deg EC, partly sloughed off; multifocal very mild pv mononuclear infiltration **vAg**: neg, apart from one bronchiole with pos intact and deg EC | 3.15e+003 |
| | | **NT (HE)**: mod mf mononuclear mucosal infiltration; focal NL infiltration and sloughed off EC and NL in lumen **vAg**: multiple disseminated patches of pos intact EC; desquamed pos EC in lumen and cell free vAg in debris in lumen | 5.79e+005 |
| 2.2 | 2 mg/kg A8 trimer, 2 h pre-infection | **Lung (HE)**: a few small consolidated areas with LC infiltration, activated and hyperplastic type II pn, several NL and numerous mononuclear cells, with desquamative component; mild mononuclearpv infiltration **vAg**: several bronchioles with patches to almost all pos BEC; rare individual pos macrophages or deg pn within inflammatory infiltrates; a few small patches of lveoli with pos AEC | 4.24e+005 |
| | | **NT (HE)**: mod mf mononuclear mucosal infiltration; sloughed off EC and NL in lumen **vAg**: multiple disseminated individual pos intact EC; desquamed pos EC in lumen and cell free vAg in debris in lumen | 1.31e+006 |
| 2.3 | 2 mg/kg A8 trimer, 2 h pre-infection | **Lung (HE)**: a few large consolidated areas with activated and hyperplastic type II pn, several NL and numerous mononuclear cells, mild desquamative component; mild pv mononuclear (LC dominated) infiltration **vAg**: neg | 6.85e+003 |
| | | **NT (HE)**: mod mf mononuclear mucosal infiltration; sloughed off EC and NL in lumen **vAg**: multiple pos intact EC; desquamed pos EC in lumen and cell free vAg in debris in lumen | 4.02e+004 |
| 2.4 | 2 mg/kg A8 trimer, 2 h pre-infection | **Lung (HE)**: extensive mf consolidation with activated and hyperplastic type II pn, several NL and numerous mononuclear cell, mild to mod desquamative component; mild to mod pv mononuclear LC dominated infiltration **vAg**: rare individual macrophages within inflammatory infiltrates; several small patches of alveoli with pos AEC | 5.15e+003 |
| | | **NT (HE)**: mod mf mononuclear mucosal infiltration; focal NL infiltration and sloughed off EC and NL in lumen **vAg**: multiple disseminated patches of pos intact EC; desquamed pos EC in lumen and cell free vAg in debris in lumen | 3.5e+005 |
| 2.5 | 2 mg/kg A8 trimer, 2 h pre-infection | **Lung (HE)**: a few large consolidated areas with activated and hyperplastic type II pn, several NL and numerous mononuclear cells, mild desquamative component; mild to mod mononuclear mononuclear LC dominated pv infiltration **vAg**: very rare individual pos macrophages or deg pn within inflammatory infiltrates; rare individual pos AEC; individual pos EC in several bronchioles and occ pos deg cells in bronchial lumen | 4.44e+002 |
| | | **NT (HE)**: mod mf mononuclear mucosal infiltration; sloughed off EC and occ NL in lumen **vAg**: multiple disseminated individual pos intact EC; desquamed pos EC and cell free vAg in debris in lumen | 8.49e+003 |
| C2.6 | 2 mg/kg A8 trimer, 2 h pre-infection | **Lung (HE)**: several small consolidated areas with activated and hyperplastic type II pn, several NL and numerous mononuclear cells, mild desquamative component; mild to mod mononuclear LC dominated pv infiltration **vAg**: individual to patches of pos BEC; numerous patches of alveoli with pos AEC and some pos macrophages within consolidated areas | 5.35e+004 |
| | | **NT (HE)**: mild to mod mf mononuclear mucosal infiltration; sloughed off EC and occ NL in lumen **vAg**: multiple disseminated individual pos intact EC; desquamed pos EC and cell free v Ag in debris in lumen | 8.65e+005 |

| | | | |
|---|---|---|---|
| 3.1 | 2 mg/kg A8 trimer,<br>24 hpi | **Lung (HE)**: several consolidated areas with activated and hyperplastic type II pn and in BEC hyperplasia, a few NL and several mononuclear cells<br>**vAg**: very rare individual pos macrophages within inflammatory infiltrates; rare pos AEC | 3.23e+003 |
| | | **NT (HE)**: very mild mf mononuclear mucosal infiltration; some sloughed off EC and debris in lumen<br>**vAg**: one individual pos intact EC; some desquamed pos EC and cell free vAg in debris in lumen | 1.76e+006 |
| 3.2 | 2 mg/kg A8 trimer,<br>24 hpi | **Lung (HE)**: several variably sized consolidated areas with activated and hyperplastic type II pn, macrophages, LC and occ NL, mild to mod desquamative component; mild to mod mononuclear LC dominated pv infiltration<br>**vAg**: rare individual posmacrophages within inflammatory infiltrates; rare small patches of alveoli with pos AEC | 7.4e+002 |
| | | **NT (HE)**: very mild mf mononuclear mucosal infiltration<br>**vAg**: neg | 3.2e+004 |
| 3.3 | 2 mg/kg A8 trimer,<br>24 hpi | **Lung (HE)**: several, partly large consolidated areas with activated and hyperplastic type II pn and BEC hyperplasia, macrophages, LC and occ NL, mild desquamative component; mild mononuclear LC dominated pv infiltration<br>**vAg**: rare individual pos macrophages within inflammatory infiltrates; rare small patches of alveoli with pos AEC; very rare pos BEC | 1.12e+003 |
| | | **NT (HE)**: (very) mild mf mononuclear mucosal infiltration, in one area (see below) loss of EC and EC in lumen<br>**vAg**: one area with a few individual pos intact EC and desquamed pos EC; cell free vAg in debris in lumen | 3.36e+005 |
| 3.4 | 2 mg/kg A8 trimer,<br>24 hpi | **Lung (HE)**: several, mainly large consolidated areas with activated and hyperplastic type II pn and BEC hyperplasia, macrophages, LC and some NL, mild desquamative component; mild to mod mononuclear pv infiltration<br>**vAg**: a few individual pos cells (macrophages and/or AEC) within consolidated areas | 5.74e+003 |
| | | **NT (HE)**: (very) mild multifocal mononuclear mucosal infiltration, in one area (see below) loss of EC and EC in lumen<br>**vAg**: one area with a few individual pos intact EC and desquamed pos EC; cell free vAg in debris in lumen | 2.69e+006 |
| 3.5 | 2 mg/kg A8 trimer,<br>24 hpi | **Lung (HE)**: one large (and one small) consolidated area with activated and hyperplastic type II pn and BEC hyperplasia, a few NL and several mononuclear cells; mild mononuclear pv infiltration<br>**vAg**: neg | 1.3e+002 |
| | | **NT (HE)**: very mild mf mononuclear mucosal infiltration; some mucus and debris in lumen<br>**vAg**: EC neg; small amount of cell free vAg in debris in lumen | 5.3e+005 |
| 3.6 | 2 mg/kg A8 trimer,<br>24 hpi | **Lung (HE)**: extensive consolidation with activated and hyperplastic type II pn, macrophages, LC and occ NL, mod desquamative component and focal with alveolar oedema; mild to mod mononuclear LC dominated pv infiltration<br>**vAg**: numerous random small patches of alveoli with pos AEC, also within areas of inflammatory infiltration | 5.25e+003 |
| | | **NT (HE)**: (very) mild mf mononuclear mucosal infiltration, in one area (see below) loss of EC and EC in lumen<br>**vAg**: one area with a few individual pos intact EC and desquamed pos EC; also cell free vAg in debris in lumen | 3.33e+005 |

| | | | |
|---|---|---|---|
| 4.1 | 0.4 mg/kg A8 trimer, 24 hpi | **Lung (HE)**: mf consolidation with activated and hyperplastic type II pn and BEC hyperplasia, mod desquamative component and several NL and numerous mononuclear cells (LC, macrophages); mild to mod mononuclear LC dominated pv and pb infiltration<br>**vAg**: rare individual pos macrophages within inflammatory infiltrates; several patches of alveoli with pos AEC, also within inflammatory infiltrates | 6.21e+003 |
| | | **NT (HE)**: very mild mf mononuclear mucosal infiltration; some mucus and debris in lumen<br>**vAg**: rare individual pos intact EC; some cell free vAg in debris in lumen | 6.16e+006 |
| 4.2 | 0.4 mg/kg A8 trimer, 24 hpi | **Lung (HE)**: mf consolidation with activated and hyperplastic type II pn and BEC hyperplasia, mod desquamative component and several NL and numerous mononuclear cells (LC, macrophages); mild mononuclear LC doinated pv and pb infiltration<br>**vAg**: rare individual pos macrophages within inflammatory infiltrates; one patch of alveoli with pos AEC | 2.33e+002 |
| | | **NT (HE)**: mild mf mononuclear mucosal infiltration; some mucus and debris in lumen<br>**vAg**: rare individual pos intact EC; some cell free vAg in debris in lumen | 2.96e+005 |
| 4.3 | 0.4 mg/kg A8 trimer, 24 hpi | **Lung (HE)**: focal areas of consolidation with activated and hyperplastic type II pn and BEC hyperplasia, marked desquamative component and several NL and numerous mononuclear cells (LC, macrophages); mild mononuclear LC dominated pv and pb infiltration<br>**vAg**: rare individual pos macrophages and AEC within inflammatory infiltrates | 4.22e+003 |
| | | **NT (HE)**: very mild mf mononuclear mucosal infiltration; some mucus with NL and debris in lumen<br>**vAg**: EC neg; small amount of cell free vAg in debris in lumen | 6.6e+005 |
| 4.4 | 0.4 mg/kg A8 trimer, 24 hpi | **Lung (HE)**: mf consolidation with activated and hyperplastic type II pn and and marked BEC hyperplasia, mod desquamative component and several NL and numerous mononuclear cells (LC, macrophages); mild mononuclear LC dominated pv and pb infiltration<br>**vAg**: rare individual pos macrophages within inflammatory infiltrates; multiple small patches of alveoli with pos AEC, mainly within inflammatory infiltrates | 5.08e+004 |
| | | **NT (HE)**: (very) mild mf mononuclear mucosal infiltration, in one area flattened epithelium, desquamed EC in lumen<br>**vAg**: one large area with several pos intact EC and desquamed pos EC; some cell free vAg in debris/mucus in lumen | 6.5e+005 |
| 4.5 | 0.4 mg/kg A8 trimer, 24 hpi | **Lung (HE)**: a few rel large consolidated areas with activated and hyperplastic type II pn, macrophages, LC and occ NL, mild desquamative component; mild mononuclear LC dominatedpv and pb infiltration<br>**vAg**: rare individual pos macrophages and AEC within inflammatory infiltrates | 2.12e+002 |
| | | **NT (HE)**: mild mf mononuclear mucosal infiltration, focally flattened epithelium; some mucus and debris in lumen<br>**vAg**: rare individual pos EC; some cell free vAg in debris in lumen | 2.86e+006 |
| 4.6 | 0.4 mg/kg A8 trimer, 24 hpi | **Lung (HE)**: small focal consolidated area with activated and hyperplastic type II pn, macrophages, LC and occ NL; (very) mild pb mononuclear LC dominated infiltration<br>**vAg**: very rare individual pos macrophages within inflammatory infiltrate | 3.67e+002 |
| | | **NT (HE)**: very mild mf mononuclear mucosal infiltration; some mucus with NL and debris in lumen<br>**vAg**: EC neg; some deg pos EC and cell free vAg in mucus in lumen | 1.15e+005 |

| | | | |
|---|---|---|---|
| 5.1 | 0.1 mg/kg A8 trimer, 24 hpi | **Lung (HE)**: extensive, often loosely consolidated areas with activated and hyperplastic type II pn, NL, macrophages and LC, mod desquamative component, occ syncytial cells, focal alveolar oedema; mild mononuclear LC dominated pv and pb infiltration **vAg**: very rare individual pos macrophages and AEC, mainly within inflammatory infiltrates | 3.29e+003 |
| | | **NT (HE)**: mild mf mononuclear mucosal infiltration, some debris in lumen **vAg**: one area with several pos intact EC, otherwise scattered individual pos EC; some cell free vAg in debris in lumen | 1.36e+007 |
| 5.2 | 0.1 mg/kg A8 trimer, 24 hpi | **Lung (HE)**: multifocal, often large consolidated areas with activated and hyperplastic type II pn and BEC hyperplasia, NL, macrophages and LC, mod desquamative component, syncytial cells, focal alveolar oedema; mild mononuclear mononuclear LC dominated pb and pv infiltration **vAg**: two small patches of pos AEC | 3.71e+003 |
| | | **NT (HE)**: mild mf mononuclear mucosal infiltration, some debris in lumen **vAg**: one area with several pos EC; some cell free vAg in debris in lumen | 5.99e+005 |
| 5.3 | 0.1 mg/kg A8 trimer, 24 hpi | **Lung (HE)**: extensive, often loosely consolidated areas with activated and hyperplastic type II pn and marked BEC hyperplasia, NL, macrophages and LC, mild desquamative component; mild mononuclear LC dominated pv and pb infiltration **vAg**: two small patches of alveoli with pos AEC and individual pos AEC, also within areas of inflammatory infiltration | 3.1e+003 |
| | | **NT (HE)**: mild mf mononuclear mucosal infiltration, focally flattened epithelium; some mucus and debris in lumen **vAg**: rare individual pos EC; some cell free vAg in debris in lumen | 1.45e+006 |
| 5.4 | 0.1 mg/kg A8 trimer, 24 hpi | **Lung (HE)**: mf, often loosely consolidated areas with activated and hyperplastic type II pn and BEC hyperplasia, NL, macrophages and LC,mod desquamative component, syncytial cells, focal alveolar oedema; mild mononuclear LC dominated pb and pv infiltration **vAg**: rare individual pos cell (macrophages, type (I and) II pneumocytes) within inflammatory infiltrates | 2.08e+002 |
| | | **NT (HE)**: mild mf mononuclear mucosal infiltration, focally flattened epithelium; abundant NL in some mucus with debris in lumen **vAg**: one pos intact EC; several pos sloughed off EC and some cell free vAg in debris in lumen | 4.53e+004 |
| 5.5 | 0.1 mg/kg A8 trimer, 24 hpi | **Lung (HE)**: extensive, often loosely consolidated areas with activated and hyperplastic type II pn and marked BEC hyperplasia, NL, macrophages and LC, mod desquamative component, focal alveolar oedema; mild mononuclear LC dominated pv and pb infiltration **vAg**: a few individual pos macrophages within inflammatory infiltrates; several, often larger random patches of alveoli with pos AEC, also within inflammatory infiltrates | 7.02e+003 |
| | | **NT (HE)**: mild mf mononuclear mucosal infiltration, abundant NL and some debris in lumen **vAg**: one area with several pos EC; some cell free vAg in debris in lumen | 1.48e+005 |
| C5.6 | 0.1 mg/kg A8 trimer, 24 hpi | **Lung (HE)**: extensive, often loosely consolidated areas with activated and hyperplastic type II pn and marked BEC hyperplasia, NL, macrophages and LC, mod desquamative component, focal alveolar oedema; mild mononuclear LC dominated pv and pb infiltration **vAg**: a few pos macrophages in inflammatory infiltrates, several random small patches of pos AEC | 2e+003 |
| | | **NT (HE)**: mild mf mononuclear mucosal infiltration, patches of mucus with embedded desquamed EC and some debris in lumen **vAg**: one area with several pos EC; pos EC and some cell free v Ag in debris in lumen | 6.31e+005 |

| | | | |
|---|---|---|---|
| 6.1 | 2 mg/kg A8 trimer, 48 hpi | **Lung (HE)**: extensive mf, mainly loose consolidation with activated and hyperplastic type II pn, NL, macrophages and LC, marked desquamative component, syncytial cells, mf alveolar oedema; bronchioles with hpyperplastic epithelium; mod mononuclear pv infiltration<br>**vAg**: several bronchioles with pos BEC; rare individual pos macrophages and AEC in inflammatory infiltrates; several disseminated small patches of alveoli with pos AEC | 3.92e+004 |
| | | **NT (HE)**: mild mf mononuclear mucosal infiltration; large NL aggregates with sloughed off EC and debris in lumen<br>**vAg**: some patches of pos intact EC; desquamed pos EC and cell free vAg in debris in lumen | 6.34e+005 |
| 6.2 | 2 mg/kg A8 trimer, 48 hpi | **Lung (HE)**: extensive mf, partly loose consolidation with activated and hyperplastic type II pn and BEC hyperplasia, NL, macrophages and LC, focal desquamative component; mild mononuclear LC dominated pb and pv infiltration<br>**vAg**: a few random individual pos macrophages and AEC within inflammatory infiltrates; a few disseminated, mainly small patches of alveoli with pos AEC | 1.33e+004 |
| | | **NT**: NA | ND |
| 6.3 | 2 mg/kg A8 trimer, 48 hpi | **Lung (HE)**: oligofocal consolidated areas with activated and hyperplastic type II pn and marked BEC hyperplasia, a few NL, more macrophages and LC, focal desquamative component, some syncytial cells; very mild pb and pv mononuclear LC dominated infiltration<br>**vAg**: very rare pos macrophages and type II pn within inflammatory infiltrates; two small patches of alveoli with pos AEC | 4.39e+002 |
| | | **NT (HE)**: mild mf mononuclear mucosal infiltration; sloughed off EC in mucus in lumen<br>**vAg**: small patches of pos intact EC; desquamed pos EC in mucus in lumen | 5.4e+005 |
| 6.4 | 2 mg/kg A8 trimer, 48 hpi | **Lung (HE)**: extensive mf consolidation with activated and hyperplastic type II pn and BEC hyperplasia, NL, macrophages and LC, mod desquamative component; mild mononuclear LC dominated pb and pv infiltration<br>**vAg**: neg | 2.34e+003 |
| | | **NT (HE)**: mild mf mononuclear mucosal infiltration; some mucus with debris in lumen<br>**vAg**: one patch of pos intact EC; small amount of cell free vAg in debris in lumen | 9.24e+005 |
| 6.5 | 2 mg/kg A8 trimer, 48 hpi | **Lung (HE)**: extensive mf, partly loose consolidation with activated and hyperplastic type II pn and BEC hyperplasia, NL, macrophages and LC, focal desquamative component; mild mononuclear LC dominated pb and pv infiltration<br>**vAg**: rare individual pos macrophages and AEC within inflammatory infiltrates; a few small patches of alveoli with pos AEC | 7.45e+003 |
| | | **NT**: very small tissue fragment, evaluation not possible | ND |
| 6.6 | 2 mg/kg A8 trimer, 48 hpi | **Lung (HE)**: a few, relatively small focal areas of consolidation, with activated and hyperplastic type II pn and a few NL and several mononuclear cells; focal mild mononuclear LC dominated pv infiltration<br>**vAg**: very rare individual pos macrophages within inflammatory infiltrates | 2.52e+002 |
| | | **NT (HE)**: very mild mf mononuclear mucosal infiltration; some mucus with debris in lumen<br>**vAg**: EC neg; small amount of cell free vAg in debris in lumen | 7.23e+004 |

**Legend**: AEC – alveolar epithelial cells; AM – alveolar macrophages; BEC – bronchiolar epithelial cells; deg – degenerate; EC – epithelial cells; HE – histological features assessed in a haematoxylin-eosin stained section; in – intranasal; LC – lymphocyte; mf – multifocal; mod – moderate; NA – not available; neg – negative; NL – neutrophils; NT – nasal turbinates; LC – lymphocytes; pb – peribronchiolar; pn – pneumocytes; pos – positive; pv – perivascular; vAg – viral antigen

[1] PCR: Copies of viral SgE-RNA/µg of RNA relative to 18S
